# Comparative Analysis of non-coding smallRNAs in *Pseudomonas aeruginosa* Keratitis Strains with Different Antibiotic Susceptibility

**DOI:** 10.1101/2021.07.02.450871

**Authors:** Kathirvel Kandasamy, Prajna Lalitha, Bharanidharan Devarajan

**Author notes:** Correspondence: Bharanidharan Devarajan, Scientist, Department of Microbiology and Bioinformatics, Aravind Medical Research Foundation, 1, Anna Nagar, Madurai, Tamil Nadu, India – 625 020.

## Abstract

Pseudomonas aeruginosa, is a gram-negative bacterium causes opportunistic or nosocomial infections in immunocompromised individuals. In recent years, a steady increase in human corneal infections of P. aeruginosa has been reported with increased multi-drug resistance (MDR) or extensively drug resistance (XDR). Several non-coding sRNAs, has been identified to regulate various physiological processes in P. aeruginosa, including biofilm formation, quorum sensing. However, the regulatory mechanism of sRNAs in MDR/XDR pathways of P. aeruginosa keratitis strains is not yet studied. In this study, we identified bacterial sRNAs in publicly available P. aeruginosa keratitis genomes and investigated their regulatory role in MDR/XDR pathways using bioinformatic analysis. Totally, 46 P. aeruginosa keratitis strains from different geographical regions were included. Of 46, Eight (30%) out of Twenty-seven and Nine (52) out of Nineteen P. aeruginosa strains from India and Australia were identified as not-MDR. Whereas, 10 (38%) Indian and 9 (47%) Australian strains were identified as MDR. Eight Indian strains were identified as XDR. Out of 46 strains, 23 (50%) carried ExoU, 21(45%) carried ExoS and two (5%) strains carried both ExoU and ExoS, exotoxins for their virulence. The sRNA, SPA0021 was identified in 18 MDR/XDR and 6 not-MDR strains along with UCBPP-PA14. Interestingly, majority of the imipenem resistant P. aeruginosa keratitis strains from the present study was found to be carried SPA0023 sRNA (18 out of 30 strains). The outer membrane porin protein OprD, identified as binding target of SPA0023. Negative regulation or inactivation of OprD, reported in increased imipenem resistance in P. aeruginosa. Mutation analysis revealed that SPA0023 carrying P. aeruginosa keratitis strains contains a lesser number of amino acid changes in OprD protein than other strains. These findings indicate, imipenem resistance in SPA0023 carried strains might arose from the negative regulation or inhibition of OprD by SPA0023. However, functional studies are warranted with large number of P. aeruginosa keratitis strains to confirm the negative regulation of OprD by SPA0023 and imipenem resistance.

## 1. Introduction

*Pseudomonas aeruginosa*, is a ubiquitous gram-negative bacterium which belongs to *Pseudomonadaceae* family that causes opportunistic or nosocomial infections in immunocompromised individuals [1,2]. *P. aeruginosa* can transform to infectious human pathogen from being an environmental isolate such as water and soil [1]. While its ability to causing chronic pulmonary inflammation in cystic fibrosis patients, urinary and respiratory tract infections is well studied, whereas *P. aeruginosa* in bacterial keratitis is relatively understudied. Human corneal infections by *P. aeruginosa* are majorly related to improper contact lens wear and handling, and other risk factors for keratitis in non-contact lens wearers includes ocular surgery, ocular trauma and earlier ocular surface disease [3-6]. Treating the infection of *P. aeruginosa* is tough due to its resistance to multiple class of antibiotics, which resulted from complex transcriptional regulatory networks it possesses and expressing different sets of genes in different environments to facilitate adaption and growth in antibiotic induced stressful environments [7,8]. In recent years, increased multi-drug resistance (MDR) or extensively drug resistance (XDR) of *P. aeruginosa* keratitis strains has been reported by several studies [9-13]. However, the emerging MDR/XDR *P. aeruginosa* keratitis strains were resistant to aminoglycosides, carbapenems and quinolones, the options for its treatment are very limited.

Bacterial non-coding smallRNAs (sRNAs) are between 50 and 300 nt long in length, induces post transcriptional events and an increased number of sRNAs has been reported in various pathogenic bacteria including *Staphylococcus aereus, Escherichia coli* and *P. aeruginosa* from the past decade [14-17]. Similar to eukaryotic miRNAs, bacterial sRNAs have multiple targets and regulate them in *trans* as well as *cis* acts by base-pairing in anti-sense orientation with complementary sequence of its target mRNAs [18]. Bacterial sRNA commonly required the sm-like RNA-binding protein called Hfq, which interacts with both sRNA and mRNA and facilitates the interaction between anti-sense sRNA and their binding target mRNAs in post transcriptionally [19,20]. Hfq also can serve alone as translational repressor of mRNA [21,22]. Several sRNAs, has been identified to regulate various physiological processes in *P. aeruginosa*, including biofilm formation, quorum sensing [17,21]. However, the regulatory mechanism of this non-coding sRNAs in MDR/XDR pathways of *P. aeruginosa* keratitis strains is not yet studied.

The aim of the present study was to identify the bacterial sRNAs in available *P. aeruginosa* keratitis genomes and study their regulatory functions in MDR/XDR pathways.

## 2. Methods

### 2.1. Bacterial strains and antibiotic susceptibility profile

In total five *P. aeruginosa* keratitis genomes from our previous study [9] and 41 reported complete genomes of *P. aeruginosa* keratitis strains from India, Australia and England [10-13] were used for non-coding sRNA identification in the present study. All genomes were retrieved in GenBank format and dated before October 2020. The details of antibiotic susceptibility of each strain used in the present study was obtained from respected reports, if available. Based on the susceptibility towards number of antimicrobial agents, the strains were defined as non-MDR, MDR and XDR [23]. Briefly, the strains which non-susceptible to ≥ 1 agent in ≥ 3 antimicrobial categories were MDR and the strains which non-susceptible to ≥ 1 agent in all but ≤ 2 antimicrobial categories were defined as XDR. The complete details of *P. aeruginosa* keratitis genomes used in this study are shown in Table 1.

**Table 1.** The complete genomic features, clinical outcome and regions of *P. aeruginosa* keratitis strains used in this study.

| Strain ID | Genome size (Mb) | No. of Contigs | CDs | GC (%) | Type | UST | Clinical Outcome | Region | GenBank accession |
| --- | --- | --- | --- | --- | --- | --- | --- | --- | --- |
| BK1 | 6.4 | 163 | 6736 | 66 | Not-MDR | ST | Poor | India | JBTQ00000000.1 |
| BK2 | 6.3 | 79 | 6037 | 66 | Not-MDR | ST | Good | India | GCA_002243265.1 |
| BK3 | 7.1 | 161 | 7009 | 66 | MDR | UT | Poor | India | GCA_002242775.1 |
| BK4 | 6.5 | 248 | 6141 | 66 | Not-MDR | ST | Poor | India | GCA_002242885.1 |
| BK5 | 6.3 | 133 | 6055 | 66 | Not-MDR | UST | Poor | India | GCA_002242915.1 |
| BK6 | 7.1 | 202 | 6722 | 66 | XDR | UT | Good | India | GCA_002242855.1 |
| PA31 | 7.1 | 137 | 6619 | 66 | XDR | UT | NA | India | GCA_003332785.1 |
| PA32 | 7.1 | 155 | 6611 | 66 | XDR | UT | NA | India | GCA_003332735.1 |
| PA33 | 7.1 | 166 | 6609 | 66 | MDR | UT | NA | India | GCA_003332715.1 |
| PA34 | 6.8 | 130 | 6326 | 66 | XDR | UT | NA | India | GCA_003332705.2 |
| PA35 | 7.1 | 156 | 6611 | 66 | MDR | UT | NA | India | GCA_003332755.1 |
| PA37 | 7.1 | 241 | 6645 | 66 | MDR | UT | NA | India | GCA_003332665.1 |
| PA82 | 6.3 | 64 | 5810 | 66 | MDR | UT | NA | India | GCA_003332645.1 |
| PA17 | 6.3 | 60 | 5825 | 66 | Not-MDR | ST | NA | Australia | GCA_003332795.1 |
| PA40 | 6.2 | 109 | 5700 | 66 | Not-MDR | ST | NA | Australia | GCA_003332655.1 |
| PA149 | 6.3 | 59 | 5745 | 66 | Not-MDR | ST | NA | Australia | GCA_003332625.1 |
| PA157 | 6.2 | 56 | 5708 | 66 | Not-MDR | ST | NA | Australia | GCA_003332575.1 |
| PA171 | 6.3 | 60 | 5812 | 66 | Not-MDR | ST | NA | Australia | GCA_003332565.1 |
| PA175 | 6.7 | 62 | 6181 | 66 | MDR | UT | NA | Australia | GCA_003332455.1 |
| PA123 | 6.3 | 86 | 5858 | 66 | Not-MDR | UT | NA | Australia | GCA_009727505.1 |
| PA126 | 6.4 | 81 | 6007 | 66 | Not-MDR | UST | NA | Australia | GCA_009727515.1 |
| PA127 | 6.3 | 86 | 5858 | 66 | MDR | UT | NA | Australia | GCA_009727535.1 |
| PA162 | 6.6 | 87 | 6131 | 66 | Not-MDR | UT | NA | Australia | GCA_009727485.1 |
| PA169 | 6.3 | 50 | 5918 | 66 | Not-MDR | UT | NA | Australia | GCA_009727465.1 |
| PA181 | 7.1 | 122 | 6619 | 66 | MDR | ST | NA | Australia | GCA_009727425.1 |
| PA182 | 6.8 | 58 | 6392 | 66 | Not-MDR | ST | NA | Australia | GCA_009727385.1 |
| PA223 | 6.9 | 108 | 6408 | 66 | MDR | ST | NA | Australia | GCA_014673185.1 |
| PA224 | 6.4 | 151 | 5811 | 66 | MDR | ST | NA | Australia | GCA_014673235.1 |
| PA225 | 7.2 | 294 | 6607 | 66 | MDR | ST | NA | Australia | GCA_014672955.1 |
| PA227 | 7.1 | 102 | 6529 | 66 | MDR | ST | NA | Australia | GCA_014673145.1 |
| PA233 | 6.3 | 92 | 5745 | 66 | MDR | UT | NA | Australia | GCA_014673245.1 |
| PA235 | 6.2 | 56 | 5719 | 66 | MDR | ST | NA | Australia | GCA_014672935.1 |
| PA188 | 6.3 | 56 | 5818 | 66 | Not-MDR | ST | NA | India | GCA_009727395.1 |
| PA189 | 6.3 | 59 | 5820 | 66 | Not-MDR | ST | NA | India | GCA_009727345.1 |
| PA193 | 6.3 | 66 | 5888 | 66 | Not-MDR | ST | NA | India | GCA_009727335.1 |
| PA198 | 7.1 | 119 | 6727 | 66 | MDR | UT | NA | India | GCA_009727325.1 |
| PA202 | 7.1 | 368 | 6883 | 66 | MDR | UT | NA | India | GCA_009727285.1 |
| PA206 | 6.5 | 55 | 6047 | 66 | Not-MDR | ST | NA | India | GCA_009727245.1 |
| PA216 | 8.3 | 1917 | 8943 | 66 | XDR | ST | NA | India | GCA_009727295.1 |
| PA217 | 6.8 | 132 | 6482 | 66 | MDR | UT | NA | India | GCA_009727225.1 |
| PA218 | 6.3 | 77 | 5840 | 66 | MDR | ST | NA | India | GCA_009727235.1 |
| PA219 | 7.4 | 166 | 7122 | 66 | XDR | UT | NA | India | GCA_009727125.1 |
| PA220 | 6.6 | 90 | 6144 | 66 | MDR | UT | NA | India | GCA_009727165.1 |
| PA221 | 7.2 | 294 | 6829 | 66 | XDR | UT | NA | India | GCA_009727135.1 |
| PA39016 | 6.8 | 486 | NA | NA | NT | UT | NA | England | AEEX00000000.1 |
| VRFPA04 | 6.8 | 1 | 5778 | NA | XDR | UT | Poor | India | GCA_000473745.1 |

### 2.2. Phylogeny and sRNA identification

A core genome based maximum-likelihood tree of 46 *P. aeruginosa* keratitis genomes was created using Parsnp version 1.2 in Harvest Suite [24]. The *P. aeruginosa* complete genomes PAO1 [1], UCBPP-PA14 [25] and taxonomic outliner PA7 [26] also included in phylogeny. Phylogenetic tree visualization and figure generation was done using the iTOL software [27]. Non-coding smallRNAs in all 46 *P. aeruginosa* keratitis genomes were identified by aligning each draft genome against Bacterial smallRNA Database (BSRD) using BLAST Ring Image Generator (BRIG) with 90% maximum and 80% minimum sequence similarity for the best hit [28]. Totally 130 known sRNAs from *P. aeruginosa* reference genome PAO1 and virulence genome UCBPP-PA14 were manually curated from BSRD for sRNA identification.

### 2.3. Target prediction and functional analysis

Bacterial binding target genes of identified sRNAs were predicted using various target prediction servers such as TargetRNA2 [29] and IntaRNA [30]. Genes with high affinity of binding with sRNAs selected for the pathway analysis using DAVID [31]and KEGG [32] databases. Gene ontology (GO) terms of identified sRNAs in Biological Process (BP), Cellular Component (CC) and Molecular Function (MF) were predicted by their binding target genes using DAVID.

## 3. Results and Discussion

### 3.1. Antibiotic susceptibility profile and genome characteristics

Totally, eighteen, nineteen and eight P. *aeruginosa* strains used in the present study were found as non-MDR, MDR and XDR respectively (Table 1). Strains with intermediate resistance to antimicrobial agents also categorized as resistant for subsequent analysis. Eight (30%) out of Twenty-six and Ten (52%) out of Nineteen *P. aeruginosa* strains from India and Australia were identified as not-MDR. Whereas, 10 (38%) Indian and 9 (47%) Australian strains were identified as MDR showing resistant to at least more than one antibiotic from ≥ 3 antimicrobial categories. Eight Indian strains were identified as XDR showing resistant to at least more than one antibiotic from all but ≤ 2 tested antimicrobial categories. Overall, Indian P. *aeruginosa* strains are more resistant to antibiotics compared to others. Among all tested antibiotics to keratitis P. *aeruginosa* strains, ceftazidime resistance in 25, ciprofloxacin resistance in 29 and imipenem resistance in 30 strains were observed (Figure 1). Strain PA193 from India showed no resistance to any tested antibiotics. All strains were sensitive to colistin. Out of 46 strains, 23 (50%) carried ExoU, 21(45%) carried ExoS and two (5%) strains carried both ExoU and ExoS, exotoxins for their virulence. Indian strain with poor clinical outcome BK5, reported to have non-synonyms mutation in ExoU protein which leads loss of ExoU gene function. Australian strain PA126 harbored both ExoU and ExoS, while no mutations were found in ExoU gene. Two not-MDR Australian strains PA126 and PA169 found to be carried ExoU. Whereas, Eight MDR/XDR strains from Australia (6) and India (2) were found to be carried ExoS (Table 1). These findings indicate ExoS carrying strains might have additional strain-specific genes for their virulence mechanism. Out of 46 strains used in this study, the clinical outcome of five strains were poor and remaining strain’s clinical outcomes were either good (healed) or not available.

**Figure 1.**
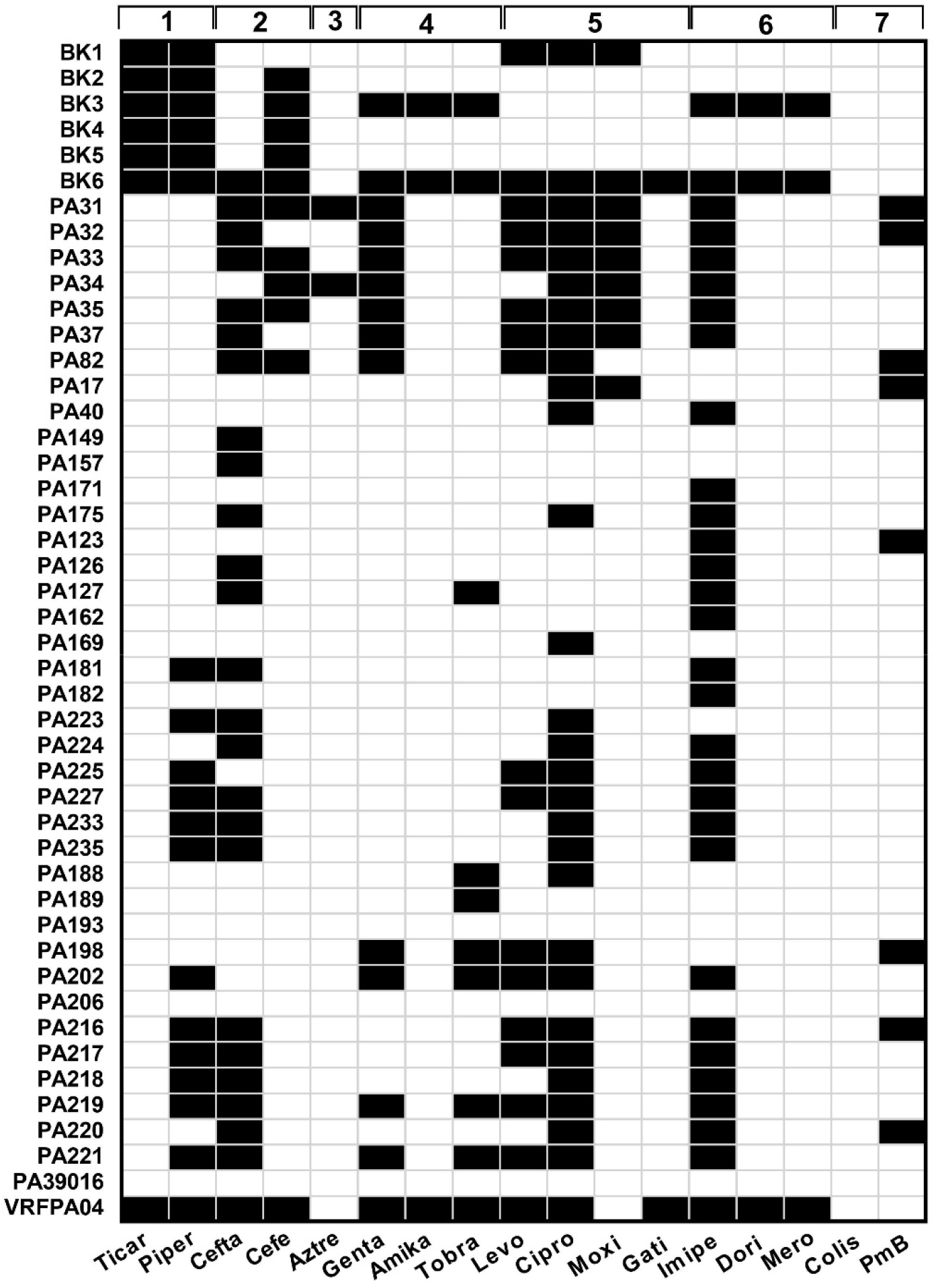
The antibiotic susceptibility profile of *P. aeruginosa* keratitis strains used in this study. Class of antimicrobial agents, [1] Penicillins/ß-lactamase inhibitors; [2] Cephalosporins; [33] Monobactams; [4] Aminoglycosides; [5] fluoroquinolones; [6] Carbapenems; [7] Polymyxins.

The genome size of keratitis *P. aeruginosa* strains ranged from 6.2Mb to 8.3Mb. The minimum and maximum number of contigs were 1 to 1917 and the average number of protein coding genes were 6285 from all *P. aeruginosa* strains used in this study (Table 1). The percentage of GC content of all strains were 66 as expected.

### 3.2. Phylogeny

Core genome based phylogenetic tree showed that all 46 *P. aeruginosa* keratitis strains were clustered into two groups, where PA7 and PA206 as outliner as expected (Figure 2). These phylogenetic results are consisted and similar with previous studies which have also shown that *P. aeruginosa* strains from different sources tend to cluster into two groups [10,34-36]. All MDR and XDR strains were clustered in group 1, along with two not-MDR strains PA162 and PA169. Group 2 contains 25 strains, where 16 and 9 were not-MDR and MDR/ XDR strains, respectively. Surprisingly, the strains were not grouped based on either antibiotic susceptibility profile or T3SS exotoxin. Group 1 tend to be smaller than group 2 and contains a greater number of Indian *P. aeruginosa* keratitis strains.

**Figure 2.**
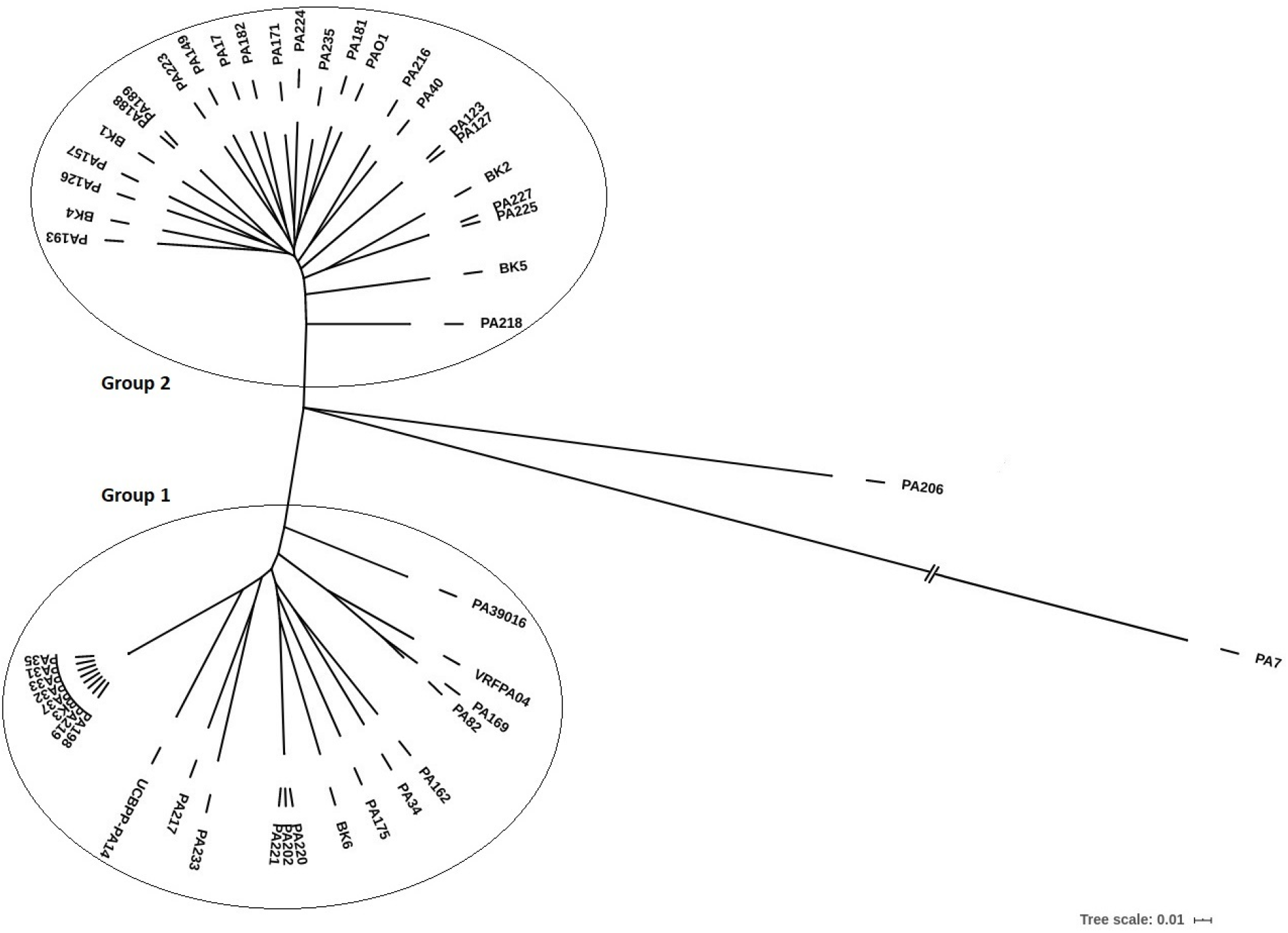
A core genome-based maximum-likelihood phylogenetic tree constructed by parsnp.

### 3.3. Bacterial non-coding sRNAs and their binding targets

We identified common and strain-specific non-coding sRNAs in *P. aeruginosa* keratitis genomes by aligning with 130 known bacterial sRNAs from PAO1 and UCBPP-PA14 reference trains using BRIG tool as shown in Figure 3. SPA0017 from UCBPP-PA14 virulent strain, was not detected or partial sequence was identified in all keratitis strains used in this study. sRNAs, SPA0010 and SPA0018 was detected only in PA39016, VRFPAO4 and UCBPP-PA14 virulent strains. The complete sequence of SPA0012 from UCBPP-PA14, was identified only in PA223 and VRFPA04. Complete sequence of UCBPP-PA14 virulent strain sRNA, SPA0011 was detected in PA175 and partially in PA223, PA225 and PA227 strains. The sRNAs, SPA0013 and SPA0019 from UCBPP-PA14 strain was not detected in any keratitis strains of the present study. Whereas, SPA0014 was detected with complete sequence identity in BK4, PA149, PA224, PA225, PA227 and VRFPAO4 strain. The sRNA, SPA0021 was identified in 6 Not-MDR and 18 MDR/XDR strains along with UCBPP-PA14. Several well-known *P. aeruginosa* sRNAs (PhrX, PhrY, PrrB, PrrH, PrrF, PrrF2 and CrcZ), reported to have role in pathogenicity was detected in all keratitis strains of the present study along with PAO1 and UCBPP-PA14 reference strains. Complete absence of PhrD was observed in all keratitis strains including UCBPP-PA14.

**Figure 3.**
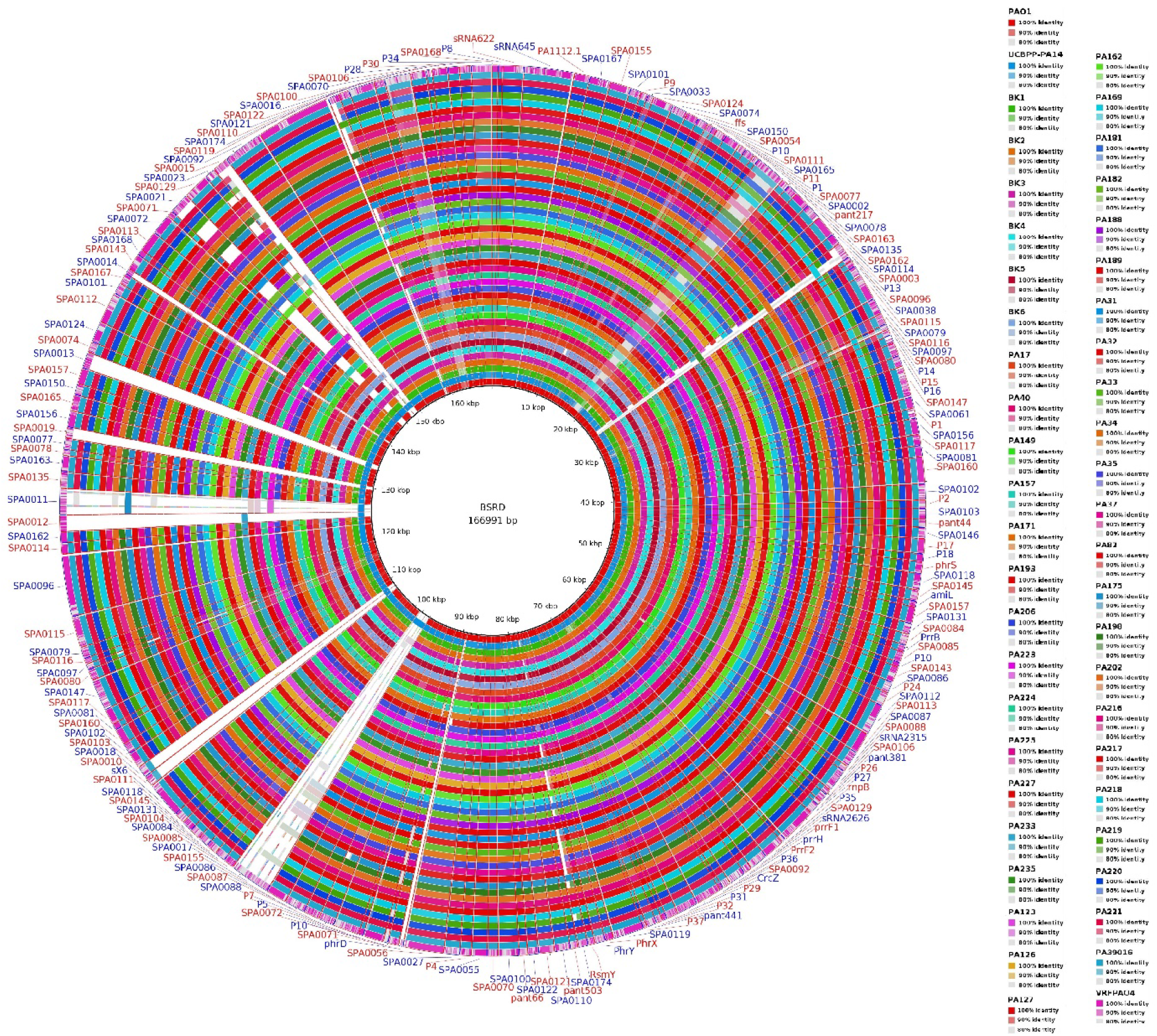
A BLAST ring circular representation of bacterial sRNAs from all *P. aeruginosa* keratitis strains used in this study.

Interestingly, sRNA SPA0023 was found in majority of the MDR/XDR keratitis strains (BK3, BK6, PA31, PA32, PA33, PA34, PA35, PA37, PA198, PA202, PA216, PA217, PA219, PA220, PA221, PA223, PA39016 and VRFPAO4) along with not-MDR strains PA162, PA182. The sRNA SPA0023, fall within the pathogenicity island PAPI-1, which is *cis*-encoded antisense to RL033 *(PA14_59840)* gene, encoding hypothetical protein. A mutation in *PA14_59840* has been reported with attenuated virulence [37]. Interestingly, majority of the imipenem resistant *P. aeruginosa* keratitis strains from the present study was found to be carried SPA0023 sRNA (18 out of 30 strains). The outer membrane porin protein OprD, identified as binding target of SPA0023 with high affinity of interaction. Negative regulation or inactivation of OprD, reported in increased imipenem resistance in *P. aeruginosa* [38-41]. Mutation analysis revealed that SPA0023 carrying *P. aeruginosa* keratitis strains contains a lesser number of amino acid changes in OprD protein than other strains. These findings indicate, imipenem resistance in SPA0023 carried strains might arose from the negative regulation or inhibition of OprD by SPA0023. However, functional studies are warranted with large number of *P. aeruginosa* keratitis strains to confirm the negative regulation of OprD by SPA0023 and imipenem resistance. The secondary minimum free energy (MFE) structure of SPA0023 was predicted using RNAfold server and shown in Supplementary Figure S1. The binding target genes of identified sRNAs were predicted by sequence-based target prediction tools TargetRNA2 and IntaRNA. Targets predicted by both tools were considered for pathway analysis and listed in Supplementary Table S1.

### 3.4. Pathway analysis

The pathways associated with binding target genes of common and uniquely shared sRNAs in keratitis *P. aeruginosa* strains were predicted using DAVID and KEGG pathway tools. Totally, 8 pathways were enriched significantly, with more than three target genes of sRNAs and listed in Table 2. The functions of predicted targets genes of sRNAs were identified in GO terms using DAVID and listed in Supplementary Table S2.

**Table 2.**
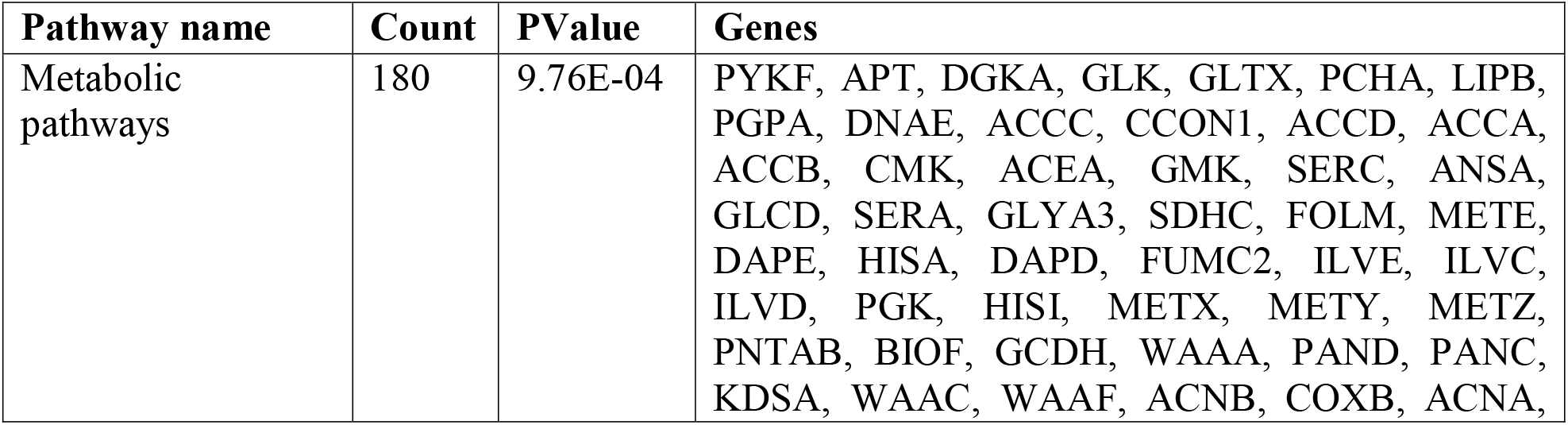
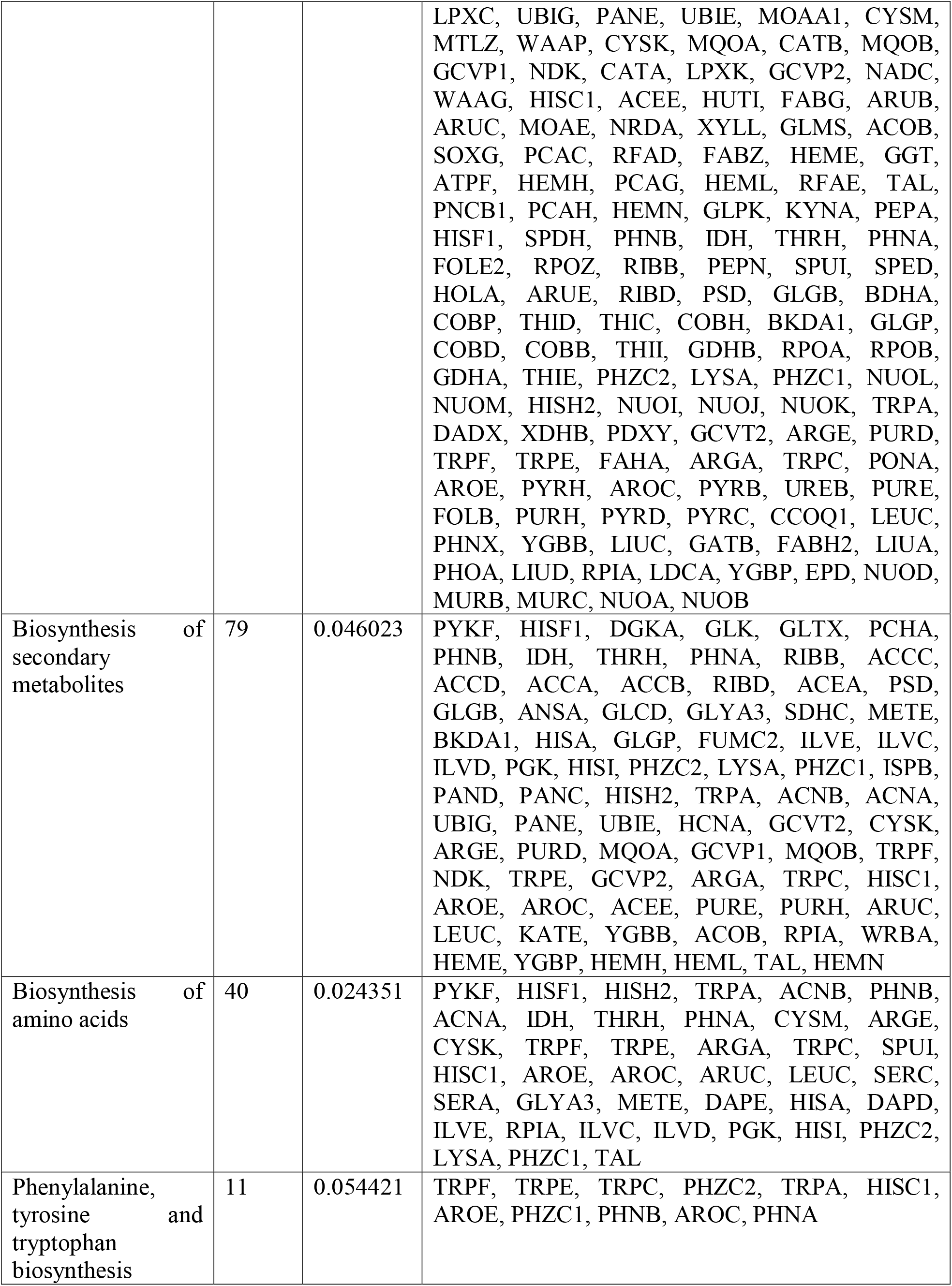
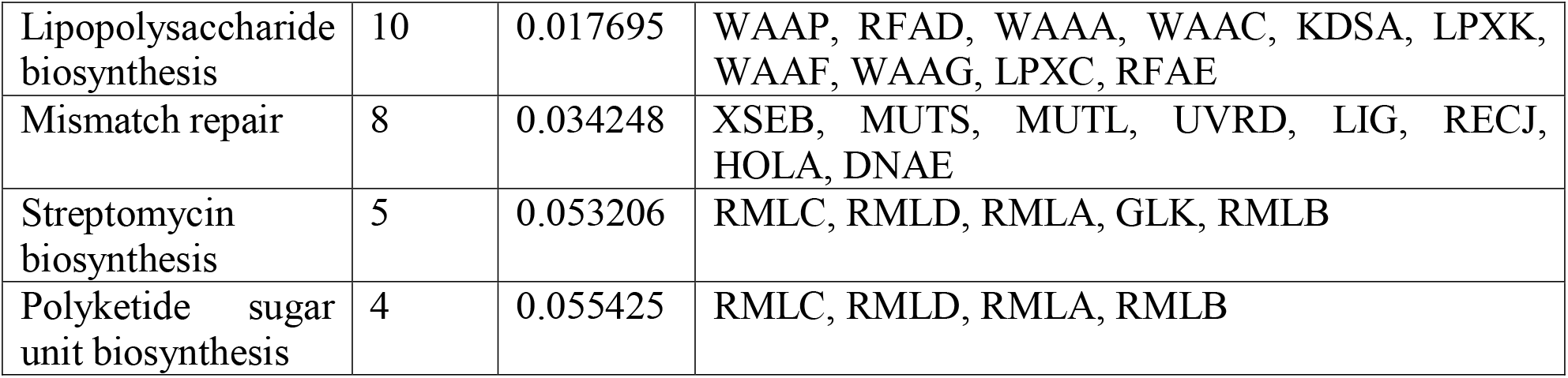
Pathways associated with target genes of identified sRNAs in this study.

## 4. Conclusion

This study identified and compared the bacterial non-coding sRNAs in *P. aeruginosa* keratitis strains with different antibiotic susceptibility profile for the first time. Totally, 46 *P. aeruginosa* keratitis strains from different geographical regions were included for the sRNA identification and investigating their regulatory role in MDR/XDR pathways. Several well-studied *P. aeruginosa* sRNAs (PhrX, PhrY, PrrB, PrrH, PrrF, PrrF2 and CrcZ), reported to have role in its pathogenicity was detected in all keratitis strains. Out of 130 known *P. aeruginosa* sRNAs, SPA0021 and SPA0023 found to expressed in majority of the MDR/XDR *P. aeruginosa* keratitis strains. Target gene prediction servers identified, outer membrane porin protein OprD was one of the binding target for SPA0023 with high binding affinity. Several studies have been reported the increased imipenem resistance with inactivation and negative regulation of OprD. However, further functional studies with greater number of *P. aeruginosa* keratitis strains are warranted to confirm the negative regulation of OprD by SPA0023.

## Competing Interest Statement

The authors have declared no competing interest.

## Funding

Aravind Medical Research Foundation, Madurai, India.

## Notes

### Summary of Updates

1. Changes in the Title made by following "Comparative Analysis of non-coding smallRNAs in Pseudomonas aeruginosa Keratitis Strains with Different Antibiotic Susceptibility" . 2. Authors list was modified into the following of " Kathirvel Kandasamy1,3, Prajna Lalitha2, Bharanidharan Devarajan1* 1Department of Microbiology and Bioinformatics, Aravind Medical Research Foundation, Madurai, India. 2Department of Microbiology, Aravind Eye Hospital, Madurai, India. 3School of Chemical and Biotechnology, SASTRA deemed to be university, Thanjavur, India." 3. The changes made in part 3.4 as followings : "Totally, 8 pathways were enriched significantly, with more than three target genes of sRNAs and listed in Table 2."

