## Supplementary Table_S1 for "Comparative Analysis of non-coding smallRNAs in *Pseudomonas aeruginosa* Keratitis Strains with Different Antibiotic Susceptibility"

**Table S1.** Identified sRNAs from *P. aeruginosa* keratitis strains and their bacterial binding target genes. Target gene prediction was done by TargetRNA2 and IntaRNA servers.

| **sRNA ID** | **Target genes** |
| --- | --- |
| amiL | coxB, ampDh3, recJ, pvdJ, phzC1, phzC2, murF, hutI, pyrB, yrfI, pslC |
| CrcZ | PA4838,PA4065,lysP,PA4066,PA2542,PA4896,PA3723,PA3632,PA1005,waaC,ribB,gdhB,prpB,arnA, waaA ,vfr,cbpA ,xcpT |
| ffs | PA0781,PA1568,rep,PA3892,PA0352,PA2224,PA2539,PA2488,opdB,PA2420,wspE,bphO,cbrA,metY, cobH |
| P1 | pslM, metK, cysS |
| P10 | fliA, vgrG1, atuR, mutM, kdpE |
| P11 | lipC, argF, trpF, phaF |
| P13 | lysA, atpS, rrmA, fliR, pscE, braE, atuR |
| P14 | IdhA, rplA, cyoA, flgI, nalD, xcpR, lysP, ponA, tauD, mutM, mexR, murA |
| P15 | PA4882,PA3858,PA2973,PA0112, chpA |
| P16 | PA1819,fabH2,lecB,PA3481,nrdJb,PA1344,amrZ,PA5182,PA1578,PA3884,hisA,trxA,hisH2,glmS |
| P17 | bkdR,PA3818,PA5062,PA3313,PA2183,PA4335,PA3390 ,PA5517,phoA,PA4882, fimL,cobB, rpmH,ftsZ, mutL,tadG,gcvT2,alkB2,aspS,psrA,glyA3 |
| P18 | rplP, ppkA |
| P2 | serA, arnA, femI, hasD, pilV, pvdR, serC, fliA, dsbB, aruC, potA, ansB, cobC |
| P24 | PA1940,ribD,PA4757,tpx, waaP, trkH,PA4859,PA2297,PA2601,cmpX ,rpsI,clpA,cyaB |
| P26 | IdhA, gcdH, iscA, ispB, hpcD, tonB1, erbR, ntrC, hemE |
| P27 | PA2116, PA3358, PA4753,PA0839,PA1881,PA2842,PA3859,PA0260,PA0309,PA2845,ftsI,  aceK, hutI |
| P28 | PA3601, liuD,PA2312,PA4961,hisJ ,PA3184,PA0797,PA3328,PA0476,PA0148, accA,pelF,algF |
| P29 | PA1891, liuA, pcaK, arnB, PA0404,PA1648,PA1593 ,phzS, rmd , mgtA,prpD,fliN , cobH,phhB, mutS, dnr,fleR, xcpQ, morB,ansA,rep,lig, iscA, fmt,mifR, fleQ |
| P30 | lig,PA5144,PA2471,PA0753,PA2529,PA2746, codB,PA3900,bioF,PA3261,dnaG,pykF,dapE,rmlC, narK2, dadX,opdH |
| P31 | carA, pro, ribC, narG, lpxK, mexD, pgpA, acoB, phzA2 |
| P32 | icp, ptsP, wzy, tadD, tag |
| P34 | thiE,PA0586,PA0586, motD,pilV,PA4841,PA4120,PA2093,PA4685,PA3170,mdcA, pcaC,phaC1, xseB,mdcC |
| P35 | gpuA, desB, modA, cpg2, bp, ggt, toxA, folM, trpA, plpD, ptxR, argS, citA, rmlC, |
| P36 | ureD, algF, liuD, parR, mtlD, glpR, katE, accA, cupA4, kynU, napA, aprF, recC, lipB, nirL |
| P37 | vfr, amiE, dht, wzm, dnaX, etfA |
| P4 | PA0203, PA2773,PA2566,PA2345 ,PA3747,PA3788 ,PA4882,PA5474,PA1012,PA0757,phzA1 |
| P5 | fliA, dsbA |
| P7 | PA4179,PA4622,PA4793,PA2336,PA2069,PA4633,metZ , senC ,PA3263,PA3390,dapE,rpoD,modC, znuB,lpxO2,tadZ,citA |
| P8 | sdsA1, qteE, ccoP2, alc, bcp, pcaH, cc4, dut, ccoP1, recC, folC, wzx |
| P9 | coIII, liuD, rpmE, fepB, leuS, hcA, cti, tpbA |
| PA1112.1 | PA4675,PA1112b,PA2212,PA3275,PA0259, PA0323, PA5059,narK2,PA1112a ,PA1037,cupA2,serA,pilX,cysS,ampDh2,znuC |
| PA1112.1 | PA4675,PA1112b,PA2212,PA3275,PA0259, PA0323, PA5059,narK2,PA1112a ,PA1037,cupA2,serA,pilX,cysS,ampDh2,znuC |
| pant217 | recG, lptF, mifS, bkdA1, ladS, pchF |
| pant44 | rhlC, tag, ureD, speA, glts, braC |
| pant441 | pmrB, ambD, ribD, aphA, fepB, motD, flgl, ptxR, treA, wspF, dbpA, creB, pilP, mifS, mdcC, dsbG |
| phrD | tolR, opmD, glpF, glnE |
| phrS | PA5318,PA0110,PA2872 ,PA2346,PA1271,PA2373,PA0071,PA4656,PA5568,PA2484 |
| PhrY | PA3961,PA1788,femI, braD ,PA3360,PA3951, rpmG, argA , atuG, hemN, cupA3 , oprP, clpB,aceE |
| PrrB | exbD1, folE2, cupA2, katB, pqL, pyrG, foxI, catA, mrcB, pilB, sbp, cupB2, hutC, pcnB, dgkA, folD, cbiD, hprA, ambD, vreA, pchG, tag, argS, glpR, aruB |
| prrF1 | katE, qscR, gabT, cc4, eraR, waaF, acnB |
| prrF2 | cc4 |
| prrH | PA2036,PA2689,PA4961,PA1103,PA3889,katE ,PA5519, qscR ,PA2173a,PA0957, gabT , cc4, cc4,waaF ,acnB |
| rnpB | PA1037,infB, gabT,PA0909,PA1243,PA1358,PA4933,PA3796,PA2852, cobD, rplR |
| SPA0002 | PA0534,PA2695,PA1467,purD,rplR ,dsdA,PA3931,PA3341,PA4713,PA2872, cmaX , rplV |
| SPA0003 | No Targets |
| SPA0010 | tpm, exsA, bacA, phnA, ylX, rhl, fxsA |
| SPA0011 | No Targets |
| SPA0012 | gloA1, ftsZ, pslH, pvdN, wspA, pvdP |
| SPA0013 | No Targets |
| SPA0014 |  |
| SPA0015 | trkA, argG, queA, hemL, atoB, recF, amiA, leuA, fruR, |
| SPA0016 | PA1407,PA1809,PA3205,PA0320, PA0367a,PA4114, cupA4 ,PA4466,PA1328,PA2556,atuF,pilV, pyrB, gabP, pcaH ,nuoK,atuH, nuoJ,pscP,aruE,fahA,gcvP1 |
| SPA0017 | No Targets |
| SPA0018 | No Targets |
| SPA0019 | spcU, nqrD, hisI, argF |
| SPA0021 | cmk, cupA2,PA3037,lysP,PA3667,icp, morB,rmlD,PA2682,PA0457,motA,fepC, pqsL, cyaB ,pyrC , gloA3 ,arnT, AmgR,mifr, dsbh,I did ,aceK |
| SPA0023 | PA1550, PA1266,oprD,pbpA,PA0698 ,PA4562,liuD, pcaR,PA4836,pcaK,argE, desA, accA ,topA,algK, tonB1,secD, ilvD,quiP,sss,hisS, pchP, pyrH |
| SPA0027 | PA2229, modA,PA1608,PA2947,PA4120,PA5168,PA0566,PA4783,PA0539,PA1894 ,erbR,phhB,tatA,pprA ,ftsZ,fmt,nalD ,arnT ,mexH ,htpX |
| SPA0033 | ntrC, exD, gln, ahpF, rpmG, aroC, nuoI, gatB, ogt, wzy, aroE, opdK, amn, flgK, purU1, liuC, pcrR, hcpC |
| SPA0038 | PA1191, PA0535,PA3754, PA1646,PA1819,ndh,PA4927,spdH ,PA3275,PA2842,micA,dhcR,thiC , rpsU,rhlI , nadC,lecB |
| SPA0054 | algF, rimM, nuoA, nusA, dsdA, Irp, pcaK, hsIU, vgrG1, rplR, xcpQ, moaA1, sdhC, algE, nqrD, mifS |
| SPA0055 | PA0355,PA0386,PA1463,PA4438,waaP , algJ, pmrB,atuR, motD , katN, creB,ptxR, gdhA |
| SPA0056 | PA2462,PA0041,PA1237, PA0570,PA4108 ,desA ,PA4836, slyD ,PA2115,PA4903, gloA3,pvcB, phnX,hisF1,potC ,dauA,mifS, vreI, truB, cupA4 |
| SPA0061 | PA4463,ssb,fumC2,PA3859, PA4150,PA0646,liuD,PA0102,wzt ,PA1419 |
| SPA0070 | PA0491,PA2074,catA ,PA2142a,PA1885,PA0384,PA3119,PA0323,PA0744 ,PA2075,selA ,vfr , metE ,acnA, phoU , fepG ,uraA |
| SPA0071 | PA0069,PA1468,ccoN1 ,morB,PA4961, PA2423, PA1492, PA2079,ispB,PA1613,argS,rplB,atpI,pdxY,ambD,prfB |
| SPA0072 | No Targets |
| SPA0074 | PA0030, aguR,PA1262,PA4887,algK,PA4181,PA4130,PA2175,bphO,PA2694, fepB, gltX, clpB, rpoD , pnp, pilH, pvcB, minC, idh, cdhA |
| SPA0077 | PA2872, femI, relA,PA2852,PA4279, PA4131,PA4219, PA4593, ercS ,PA4377, cysK ,rmlD |
| SPA0078 | desB,tpx,cynR,PA1417,PA2601, argS, pslM , rpoZ,PA4488,PA4983,priA,mexD ,xseB, phzC2 ,wspD ,lipH , phzC1, pgk |
| SPA0079 | PA0637,PA4838,PA3754, PA1819, PA4041,PA3526, PA0221 ,PA1646,PA2062,PA1390, obg,cpo |
| SPA0080 | PA2723,PA2842,PA0479,PA1819,PA2187,folB,PA4282,PA1035, PA3754,PA5168, exsE ,atuD, ptxR,mifR, phzD1,phzD2,migA,wspD,dadA |
| SPA0081 | PA3069,PA2699, PA3173, PA4070,PA4824, smpB ,motD ,PA5270,PA5254,PA3326, nuoM |
| SPA0084 | PA3764,PA1819,PA5473 ,PA4682,PA3313 ,PA1646,PA4882,PA1700, pqsC ,PA4929 |
| SPA0085 | PA3175,PA3492,thiI,PA2795,PA1361,PA4987, hpaA ,PA5212,PA0697,PA3053,pyrC,sbcD,nalD,cupA2,ptrB, glpF, sodM,xseB,dadX,nosY,bdhA,nosL,uvrD,mqoB, gloA3,pchG |
| SPA0086 | PA4793, amgR,PA0014, phuR,PA4390, sodM,PA4684,PA1880,PA2795,PA3489,ureD , xseB,lnt,wbpY, desB ,trxB2 ,argS, braD,pcaK ,tesB,mtlD, ubiG |
| SPA0087 | PA4100, PA4757, PA4169, purH , rmlB ,PA1646, kynA,PA1762, cpg2, pslM |
| SPA0088 | recF, PA3754, PA3313,cobP, PA4336, PA1878,PA5528, PA4539, exsA ,PA1699, fklB |
| SPA0092 | PA4838,PA4065,lysP,PA4066,PA2542,PA4896,PA3723,PA3632,PA1005,waaC,ribB,gdhB,prpB,arnA, waaA ,vfr,cbpA ,xcpT |
| SPA0096 | PA4420,PA3913,PA0742, mqoB, nuoL, oprD,PA2972, PA5158,PA4860, cynT,cpo, pepN,waaL,pncB1 |
| SPA0097 | pilK, thrH, alkB2, lpxC, ambD, yl, pcaG, fliN, argA, murC |
| SPA0100 | PA1819, metZ, cupB4, ssb,PA0634,PA0838,PA0588,bfrA,PA4491,PA1103,atuB,fpvB, popD , lipB,moaE ,ftsA,def, phzC1,flhA, napD,phzC2 ,rcpA ,fis,pchC |
| SPA0101 | PA1244,PA4595,PA5266, PA4824 , PA1361,PA3595,PA3973,PA5075,PA2027,PA1089, lpxK, amgR, cysM |
| SPA0102 | PA3576,PA2571,PA3462, rpsA,PA3390,PA3632 ,PA3660, PA1103,PA2568,PA0343,pepA , argS , rpsO |
| SPA0103 | femI,PA2366,fadH1,PA0940, PA2791, PA0093,PA3979, PA0218, PA3249,PA0389,creC |
| SPA0104 | rhlI,ygbP , rhlR ,aruB ,lpxO1, femA, exsA, pslM,nqrA,rhlA,ilvE ccmA ,atpF, lepB,hemK |
| SPA0106 | PA5462,PA4984,ureD,PA4746,PA4191,apt, PA1026, PA2718,PA2055,PA2714,rpsR , glyQ,flgL,xdhB, motD, trpC |
| SPA0110 | sodM,PA0875,PA2896,PA0924,braD, oprL, PA0334,PA1961,PA4340,PA0272, vqsM, aceA,recD, psd,lepA |
| SPA0111 | PA3949, cmk,PA2408, PA1315, PA1608,PA4983,PA4038, PA4145,PA5103,PA2933, pctC, aruC, phuR, wspA ,dgkA,trpE, pvdO,dnr,spuI,pchP |
| SPA0112 | PA4066,PA1827 ,ptxR,PA4131,PA4282,PA5474,PA0182,PA5436, typA ,PA4869, pslC , trpC |
| SPA0113 | PA1262,PA4466, PA4637a,PA0824,recJ ,PA2654,PA2165,PA3848,PA4093,erbR,anr,uvrA ,xcpY, pilX , glcD, hisS ,pslC,fliP , fabZ, htpX |
| SPA0114 | PA1977,PA1290 ,PA2723 , PA3081,PA3313,PA3047, PA4900,PA1652,PA3127,PA4838, mutL,prlC,soxG,glgP,ccoN1,nuoJ,dauA,pscU ,pvcC,rmlA |
| SPA0115 | PA2261,fabZ ,PA0022,sodM,PA3754,PA3969a,PA1261,PA0709,PA1831,PA0487,phhB,xseB ,dsdA, rcpA,mliC, ccoN1,gloA3 , rubA2,braD,fliJ |
| SPA0116 | PA5471,PA4596,PA1824,PA1807 ,PA0150,PA0627,PA2026,PA3325, phoB, PA4753,gltS ,ureB , hemL, rnc , accC ,pscU |
| SPA0117 | PA2154, PA5252a, ygbB,PA2123,PA5330, PA1286,PA5486, murB ,PA4186,PA0218,ftsI,uvrA |
| SPA0118 | PA1050, fliQ,PA2772,PA1237, PA1810,PA3670,PA0812,PA1243, lepA,PA2362,cueR, wbpL, rpiA ,lysA |
| SPA0119 | PA2534,PA1287,PA3085,PA2561,PA1626,PA2571,PA1164, panD, rplV,PA3667,hutI ,pvdL,rfaE,osmC , purE,xcpT |
| SPA0121 | PA0685, oprE,PA2230,metX ,PA0246,PA1849, arnB ,PA5390, exsC,PA1749, accD , micA,dnaE ,phhB , hflC |
| SPA0122 | PA3949, PA0205,PA1147,PA1361, PA1315,ptsN,PA4149,icp,PA1966,epd,recR,bcp,pilB,dapE,motA,cmk,coxB, gmk,pyrC,nosY, moaA1, aruC,metR ,moeA1, argS ,panE,metY , accB, dadX,dgkA, fepB,tpbA,mqoB,moaB1,grpE, cupC3 ,arnB,oprL,lptF |
| SPA0124 | PA2791,PA3022, dapD , nqrA ,PA4171,PA2077,PA1256,PA1764,PA1388,PA5273 |
| SPA0129 | PA2947, PA1254,PA3788, PA2455,PA4118, PA3461,PA4894,pepN,mutM,wrbA,phnB ,panC,pntAB,ribB , pncB1 |
| SPA0131 | PA4798,PA0361, PA0394,PA3676,PA5023,rpmJ,PA0781,PA2868, pyrD,PA1111,sss,desT, mutM, nrdA,dsbG,ilvC ,ftsJ,mtlZ |
| SPA0135 | PA3220,PA3510,PA1026,thiE,PA2435, xcpQ, wapR,PA2559,PA2099,PA2833, motD,hscA,amiA, mltB1 ,pmrB,pscU |
| SPA0143 | PA5246, ldcA,PA2182,PA2362,PA3252, purD,PA2116,PA1607, recN , nuoB |
| SPA0145 | argE, PA2363,PA0011,PA3474,PA3275,PA1579,PA1875, mqoA ,PA0246 |
| SPA0146 | pvdP,PA4606,PA5472,PA0452 ,PA1759,PA5003,PA4352,PA1932,PA2548,PA4337 , pgpA, catB, pcaK ,phnA ,rph |
| SPA0147 | rluC,PA0902,PA1256,rep,PA2880,PA3465,tadG ,PA2098, phzD1, phzD2, oprM,napB ,ileS,tyrS,argA,waaG,fmt , hemN ,hcnA,pys2, groES,thiD |
| SPA0150 | pyrD,PA3205,PA3526,PA2163,PA3324,pchD,PA2440 ,PA0058, xseB,PA1297, creD,rplW,nosY , ampC |
| SPA0155 | PA2842, PA1655,PA0219, xcpY ,PA0234,tal, ccoQ1,cyoE,lasB,pilW |
| SPA0156 | rplF, PA2822,PA1550,wzm,PA3510,PA1065,PA5101,PA0726, pmpR , PA2954 |
| SPA0157 | flgE,PA2654, PA4677,cdhA,hisC1,kdsA, PA3334, creD,PA5484,PA0343,bphO, leuS ,glnE |
| SPA0160 | PA4145,ldhA,coxB ,PA3939,PA5486, folE2 ,PA5127,PA3265,PA3502 ,uvrB ,rfaD,rpsI |
| SPA0162 | PA0880, hslU,PA1977,PA1262,glcD,PA1629,PA2597,PA2791,PA1465,PA1993,dhcA,braE,bdhA,pscI,rbsR **,** mobA,cspD, mifS,mifR |
| SPA0163 | PA0491,PA2569 ,PA2858,PA5382,rep,PA1613, PA5185, metE , rpoZ, trpF ,nuoL,desB , lysP ,gyrA, vfr,gpuA |
| SPA0165 | PA4420,PA5024,serC,PA4950,PA5210, cobH ,catA,PA3124,PA2722,nosY, truA,hisI,soxG,arsC,xylL, atuE |
| SPA0167 | ubiB,PA0733,fliG, pcpS,pchA,PA0570, PA2018,PA0093,PA2230 , gcvP2 |
| SPA0168 | mexE, glgB, ftsI, vreI, alkB2 |
| SPA0174 | PA2922,PA1879,PA2778,PA0571, pqsH ,mtlZ ,PA1030,PA3733a,rnt,PA4548 , wrbA |
| sRNA2315 | phzE1, phzE2, epd, pyrC, sbcD, dadX, bcpp, nalD, yfiR, fabG, recJ, eco, ptsP, ptxR, cbrA, accB, nosY, xseB, napD, liuC, hemE, typA, cmk, mqoB, |
| sRNA2626 | infB, ldcA, ptsN, zipA, pqqA, map, pslM, rpoB |
| sRNA622 | PA3538,PA0133, alkB1,femI ,PA5402,PA4968 ,fliN, gidB, smpB,PA1057,glgP ,rpoA ,uraA |
| sRNA645 | hpaA,PA1367, PA1112a,PA4987 , tag,PA5566,PA4149,PA1541,PA2650,PA5136,nosY,glpK, narK2,nalD,phaD , bkdA1,serA,antA,cysS ,pcrD ,xseB ,serC ,erbR |
| sX6 | leuC,surA , spuF,PA2408,PA0383 ,PA0475 ,oprD, infB ,pcaQ ,trpC ,ldcA ,ponA,phaF, czcB , pscD,xseB,acoB |
| PhrX | glt, ubiE, aceK, xcpQ, hemH, fabG, noY |
| PhrY | femI, braD, rmpG, argA, atuG, hemN, cupA3, oprP, clpB, aceE |
| pant503 | aguR, nirB, vqsM, hisI, pcaF, ndk, dsbA, dhcB, hflK, IdhA |
| pant66 | metZ, glk, flhA, rcpA, kdpC, bdhA |
| pant381 | morB, acoB, tyr, prpD, aguB, argG, ureB, mutM, metY, gyrA, dauA, bphO, mexG |
| RsmY | epd, aceE, coaD, masA, tatC, mvaT, xerD, cpG, trkH, flgC, moeA1, napA, treA, gatA, kefB, glpK, trpS, |
