## Supplementary Table_S2 for "Comparative Analysis of non-coding smallRNAs in *Pseudomonas aeruginosa* Keratitis Strains with Different Antibiotic Susceptibility"

**Table S2.** Gene Ontology of identified sRNAs from this study. The functions of sRNAs were identified using DAVID pathway analysis tool, BP - Biological Process; CC - Cellular Component; MF – Molecular Function.

| **Category** | **Term** | **Count** | **Fold Enrichment** | **PValue** | **Genes** |
| --- | --- | --- | --- | --- | --- |
| BP | phenazine biosynthetic process | 9 | 3.56 | 0.001 | PHZE2, PHZD1, PHZE1, PHZC2, PHZC1, PHNB, PHZD2, PHZA1, PHNA |
| BP | lipopolysaccharide core region biosynthetic process | 7 | 3.17 | 0.014 | WAAP, RFAD, WAAC, LPXK, WAAF, WAAL, RFAE |
| BP | tricarboxylic acid cycle | 9 | 2.59 | 0.014 | ACEA, MQOA, MQOB, ACEK, ACNB, SDHC, ACNA, IDH, FUMC2 |
| BP | tryptophan biosynthetic process | 6 | 3.17 | 0.028 | TRPF, TRPE, TRPC, TRPA, PHNB, PHNA |
| BP | quorum sensing | 6 | 3.17 | 0.028 | PQSH, RHLI, QSCR, RHLR, VQSM, QUIP |
| BP | positive regulation of transcription, DNA-templated | 7 | 2.61 | 0.038 | CUER, PCAR, EXSC, NOSR, PCAQ, PSRA, PTXR |
| BP | translation | 17 | 1.63 | 0.045 | RPLR, DEF, RPLP, RPSU, LEPA, GATB, RPSA, RPLV, RPLW, TYPA, RPSI, PRFB, RPSR, RPSO, RPMJ, RPMG, RPMH |
| BP | lipid A biosynthetic process | 7 | 2.46 | 0.050 | WAAA, FABZ, LPXK, ARNT, ARNA, ARNB, LPXC |
| BP | cellular amino acid biosynthetic process | 8 | 2.20 | 0.056 | CYSK, TRPF, TRPE, TRPC, TRPA, AROE, AROC, CYSM |
| BP | enterobactin biosynthetic process | 3 | 6.34 | 0.066 | PCPS, PHZD1, PHZD2 |
| BP | malonyl-CoA biosynthetic process | 3 | 6.34 | 0.066 | ACCC, ACCD, ACCA |
| BP | fatty acid biosynthetic process | 8 | 2.11 | 0.070 | FABZ, PQSC, ATUH, FABH2, ACCC, ACCD, ACCA, ACCB |
| BP | tetrahydrofolate biosynthetic process | 5 | 2.88 | 0.079 | PHZE2, PHZE1, THRH, FOLB, FOLE2 |
| BP | pilus assembly | 4 | 3.62 | 0.082 | CUPA3, CUPC3, PILB, WAAL |
| BP | lipopolysaccharide biosynthetic process | 10 | 1.76 | 0.099 | KDSA, RMLC, WZY, WAAF, RMD, ARNT, ARNA, ARNB, WBPY, WBPL |
| CC | cytoplasm | 102 | 1.43 | 1.98E-05 | HISS, SLYD, CLPB, APT, CDHA, GLK, CHPA, PCHA, PILB, LIPB, DNAE, PNP, MDCC, PRFB, RUBA2, CMK, ACEA, GMK, SERC, GLYA3, HISA, DAPD, MOAB1, PGK, HISI, METX, METY, METZ, PAND, PANC, KDSA, MOBA, ACNA, UBIG, UBIE, CYSM, PRPB, MOEA1, WAAP, CYSK, NDK, ACEK, NADC, HUTI, SELA, EXSC, DSDA, FABZ, HEMH, PTSN, CSPD, OGT, HEML, TAL, HEMN, GYRA, CUER, GLPK, INFB, PEPA, HISF1, RNC, TRXB2, FOLE2, RIBB, FTSZ, PEPN, TPBA, ASPS, GLGP, UVRD, RPOD, UVRB, UVRA, THIE, TRXA, GIDB, ARGS, HISH2, TRPA, ARGE, ARGA, OBG, FKLB, URED, PYRB, UREB, PYRD, PHZE2, PHNX, PHZE1, FABH2, SMPB, PHOB, RECF, XSEB, SSS, LDCA, EPD, PHOU, MURB, MURC |
| CC | motile cilium | 7 | 2.86 | 0.026 | FLGE, MOTD, FLIJ, FLGK, FLGL, FLIQ, FLHA |
| CC | cytosol | 51 | 1.30 | 0.030 | GRPE, FTSJ, BKDR, GLTX, PHNB, TOPA, ATUR, PMPR, GLYQ, RIBD, TYRS, THIC, ISCA, METE, DAPE, FUMC2, ILVE, ILVD, THII, ILES, DEST, WAAC, PHZD1, WAAF, ACNB, CYSS, PANE, XDHB, NALD, PDXY, GCVP1, GCVP2, PSRA, PHZD2, PYRH, AROC, PRLC, AGUR, ARUC, MOAE, KATE, GLMS, LIG, DHCR, RLUC, NUSA, HEME, LEUS, MORB, GLNE, VQSM |
| CC | anthranilate synthase complex | 3 | 5.52 | 0.091 | TRPE, PHNB, PHNA |
| CC | acetyl-CoA carboxylase complex | 3 | 5.52 | 0.091 | ACCD, ACCA, ACCB |
| MF | transferase activity, transferring glycosyl groups | 6 | 5.48 | 0.001 | MIGA, WAPR, WAAF, PONA, WBPL, PSLC |
| MF | quinone binding | 8 | 3.24 | 0.006 | NUOL, NUOM, NUOI, NUOJ, NUOK, NUOD, NUOA, NUOB |
| MF | NADH dehydrogenase (ubiquinone) activity | 7 | 3.65 | 0.007 | NUOL, NUOM, NUOJ, NUOK, NUOD, NUOA, NUOB |
| MF | cysteine synthase activity | 4 | 7.31 | 0.009 | CYSK, METY, METZ, CYSM |
| MF | zinc ion binding | 24 | 1.70 | 0.009 | HUTI, CYNT, MUTM, CPG2, GLTX, CYSS, THIC, METE, DAPE, FOLE2, GLOA3, DNAG, PTRB, ARGE, ILES, PEPN, HISI, HTPX, UVRA, ACCD, RIBD, PYRC, PRIA, ARUE |
| MF | acetyl-CoA carboxylase activity | 4 | 5.84 | 0.020 | ACCC, ACCD, ACCA, ACCB |
| MF | transcription regulatory region sequence-specific DNA binding | 5 | 4.06 | 0.024 | PSRA, AGUR, DEST, ATUR, NALD |
| MF | ATP binding | 73 | 1.25 | 0.024 | HISS, GYRA, GLPK, MUTS, MUTL, CLPB, CLPA, MODC, DGKA, GLK, SPUF, HSCA, GLTX, PPRA, PILB, ATUF, GLYQ, ACCC, ACCD, ACCA, CMK, TYRS, PVDE, GMK, COBP, THID, ASPS, KDPC, COBB, THII, ILES, UVRD, PGK, UVRB, UVRA, PRIA, UBIB, ARGS, PANC, PMRB, CYSS, MIFR, MIFS, WZT, PDXY, WAAP, CBRA, CREC, PURD, FLEQ, FEPC, NTRC, FLER, NDK, LPXK, ACEK, PYRH, NRDA, ZNUC, GATB, CCMA, LIUD, RECF, RECD, PPKA, LEUS, HSLU, RECN, GROES, REP, MURC, GLNE, RFAE |
| MF | structural constituent of ribosome | 14 | 1.82 | 0.034 | RPLR, RPLP, RPSU, RPSA, RPLV, RPLW, RPSI, RPLB, RPSR, RPSO, RPMJ, RPLF, RPMG, RPMH |
| MF | single-stranded DNA binding | 4 | 4.87 | 0.036 | RECF, SSB, MUTL, REP |
| MF | pyridoxal phosphate binding | 16 | 1.69 | 0.042 | BIOF, ARUC, SERC, TRPA, GLYA3, DSDA, DADX, CYSM, GLGP, ILVE, CYSK, LYSA, HISC1, METY, METZ, HEML |
| MF | ferric iron binding | 4 | 3.65 | 0.083 | CATA, BFRA, PCAG, PCAH |
| MF | anthranilate synthase activity | 3 | 5.48 | 0.092 | TRPE, PHNB, PHNA |
