## Supplementary figures and images for "Comparative Analysis of non-coding smallRNAs in *Pseudomonas aeruginosa* Keratitis Strains with Different Antibiotic Susceptibility"

### Supplementary Figure_S1

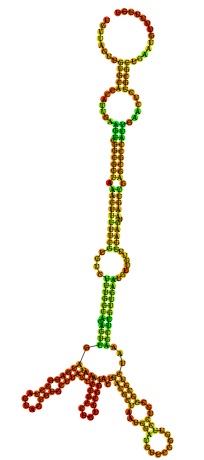
